# *TAB1* and *ASP1* act antagonistically on cytokinin signaling to regulate axillary meristem formation in rice

**DOI:** 10.64898/2026.05.19.726093

**Authors:** Ami Ohyama, Taiyo Toriba, Moeko Sato, Hiroyuki Tsuji, Wakana Tanaka

**Affiliations:** Graduate School of Integrated Sciences for Life, Hiroshima University, 1-4-4 Kagamiyama, Higashi-Hiroshima, Hiroshima, 739-8528 Japan; School of Food Industrial Sciences, Miyagi University, 2-2-1 Hatatate, Taihaku-ku, Sendai, Miyagi, 982-0215 Japan; Bioscience and Biotechnology Center, Nagoya University, Nagoya, Aichi 464-8601, Japan; Kihara Institute for Biological Research, Yokohama City University, Maioka 641-12, Totsuka, Yokohama 244-0813, Japan

## Abstract

Plants continuously develop shoot branches derived from axillary meristems. In rice (*Oryza sativa*), *TILLERS ABSENT1* (*TAB1*), an ortholog of Arabidopsis *WUSCHEL*, plays an essential role in axillary meristem formation by promoting stem cell proliferation. Although several genes associated with *TAB1* function have been identified, the molecular mechanisms underlying stem cell proliferation during axillary meristem formation remain poorly understood. Here we identify ABERRANT SPIKELET AND PANICLE1 (ASP1), a TOPLESS-like transcriptional corepressor, as a novel regulator of axillary meristem formation, and investigate downstream mechanisms regulated by TAB1 and ASP1. In *asp1*, the stem cell region was expanded, indicating that *ASP1* negatively regulates stem cell proliferation. Notably, *WOX4*, a paralog of *TAB1*, was precociously expressed in *asp1*, possibly in association with expansion of the stem cell region. Genetic analysis further revealed that *asp1* mutation rescued the loss of axillary meristems in *tab1*. Transcriptome analysis showed that several type-A *RESPONSE REGULATOR* (*OsRR*) genes, encoding negative regulators of cytokinin signaling, were upregulated in *tab1* relative to wild type, *asp1*, and the *tab1 asp1* double mutant. Consistently, fluorescence of the synthetic cytokinin reporter was absent during axillary meristem formation in *tab1* but was detected in wild type and *tab1 asp1*. Moreover, overexpression of *OsRR10* inhibited axillary meristem formation, phenocopying *tab1*. Collectively, these findings suggest that TAB1 activates cytokinin signaling by repressing type-A *OsRR* expression, whereas ASP1 negatively regulates cytokinin signaling by promoting the expression of these genes. Thus, rescue of the *tab1* phenotype by *asp1* mutation probably reflects restoration of cytokinin signaling.

## Introduction

The development of shoot branches is a key determinant of plant architecture and an important trait related to the yield of many crops. This developmental process can be broadly divided into two phases. The first phase is axillary meristem formation, during which an axillary bud containing the axillary meristem harboring stem cells is produced at the boundary between the stem and leaf. In the second phase, referred to axillary bud outgrowth, the axillary bud is released from dormancy to allow the axillary meristem to actively produce leaves and stems, eventually giving rise to a shoot branch. While extensive progress has been made in understanding the mechanisms regulating axillary bud outgrowth, those underlying the first phase of axillary meristem formation remain poorly understood (Barbier et al. 2019; Tanaka et al. 2023; Yang et al. 2023).

In grasses including rice (*Oryza sativa*), shoot branches—also referred to as tillers—are important for grain yield and biomass production. Axillary meristem formation in rice involves dynamic morphological changes that are distinct from those in eudicots such as Arabidopsis (*Arabidopsis thaliana*). It begins in the axils of the plastochron two (P2) leaf of the main stem, where *OSH1*, a marker of undifferentiated cells, is expressed (Oikawa and Kyozuka 2009; Tanaka et al. 2019) (Supplemental Fig. S1). Key regulators of this initial stage are *MONOCULM1*, *LAX PANICLE1* (*LAX1*), and *LAX2* (Komatsu et al. 2003; Li et al. 2003; Oikawa and Kyozuka 2009; Tabuchi et al. 2011). Subsequently, a small bulge termed the ‘pre-meristem zone’ emerges at the axils of P3 or P4 leaves, where axillary stem cells are established (Tanaka et al. 2015; Hirano and Tanaka 2020; Tanaka and Hirano 2020) (Supplemental Fig. S1).

In rice, *TILLERS ABSENT1* (*TAB1*), an ortholog of Arabidopsis *WUSCHEL* (*WUS*) belonging to the WOX transcription factor family, is transiently expressed in the pre-meristem zone, where it is required for stem cell proliferation (Tanaka et al. 2015; Tanaka and Hirano 2020; Ohyama and Tanaka 2025). In a loss-of-function mutant of *TAB1*, stem cell proliferation in the pre-meristem zone is impaired, resulting in defective axillary meristem formation and loss of tillers. The expression of *TAB1* is negatively regulated by *FLORAL ORGAN NUMBER2* (*FON2*), an ortholog of Arabidopsis *CLAVATA3* (*CLV3*) encoding a CLE peptide precursor (Suzaki et al. 2006; Tanaka and Hirano 2020). In a loss-of-function mutant of *FON2*, the expression domain of *TAB1* is expanded in the pre-meristem zone, leading to the overproliferation of stem cells (Tanaka and Hirano 2020). Thus, the FON2–TAB1 pathway plays a central role in maintaining proper stem cell numbers in the pre-meristem zone during axillary meristem formation in rice. At a later stage, a dome-shaped meristem emerges (Supplemental Fig. S1) and *TAB1* expression ceases; instead, its closest paralog, *WOX4*, is expressed, promoting stem cell proliferation in the developing meristem (Ohmori et al. 2013; Tanaka et al. 2015). Therefore, two WOX genes (*TAB1* and *WOX4*) act sequentially as positive regulators of stem cell proliferation during axillary meristem formation in rice.

In contrast to rice *TAB1*, which functions only within a narrow developmental window, Arabidopsis *WUS* positively regulates stem cell proliferation across all aerial meristems, including the shoot apical meristem (SAM) and the axillary meristem (Laux et al. 1996; Mayer et al. 1998; Schoof et al. 2000; Wang et al. 2017). The role of *WUS* is well characterized, particularly its close association with the plant hormone cytokinin, which activates *WUS* expression (Gordon et al. 2009). This expression is induced by type-B Arabidopsis RESPONSE REGULATORs (ARR1, ARR2, ARR10–ARR12)—positive regulators of cytokinin signaling that directly bind to the *WUS* promoter (Meng et al. 2017; Wang et al. 2017). Conversely, WUS directly represses the transcription of type-A Response Regulators (ARR5–ARR7, ARR15), which are negative regulators of cytokinin signaling, leading to activation of the cytokinin signaling pathway (Leibfried et al. 2005; Zhao et al. 2010). This positive feedback loop between WUS and cytokinin signaling is crucial for maintaining stem cell homeostasis in meristems. Despite extensive knowledge of WUS function in Arabidopsis, however, the factors regulated by TAB1 to promote stem cell proliferation during axillary meristem formation in rice remain largely unknown.

*TOPLESS* (*TPL*) and *TOPLESS-RELATED* (*TPR*), encoding transcriptional corepressors of the Groucho/Tup1 family, were first identified for their roles in apical–basal patterning during embryogenesis in Arabidopsis (Long et al. 2002; Long et al. 2006). In a gain-of-function dominant-negative *tpl-1* mutant and multiple *tpl*/*tpr* loss-of-function mutants, roots develop in place of shoots, indicating that *TPL* and *TPR* regulate shoot fate determination during embryogenesis. TPL and TPR are critical factors in several plant hormone signaling pathways, functioning not only in embryogenesis but also in other developmental processes including meristem maintenance (Liu and Karmarkar 2008; Plant et al. 2021). Among their known interaction partners is WUS (Kieffer et al. 2006); and in Arabidopsis, the TPL–WUS interaction has been implicated in the maintenance of reproductive meristems (Causier et al. 2012). Notably, *tpl tpr1 tpr4* mutants exhibit defects in axillary meristem formation, indicating that *TPL* and *TPR* are also involved in this process (Liu et al. 2024).

In maize (*Zea mays*), the *TPL* and *TPR* homologs *RAMOSA1 ENHANCER LOCUS2* (*REL2*) and *REL2-LIKE* (*RELK*) have well-characterized roles in meristem regulation. The *rel2* mutation was originally identified as an enhancer of the highly branched inflorescence phenotype of *ramosa1* (Gallavotti et al. 2010). Both *rel2* and *rel2 relk1* mutants develop an enlarged inflorescence meristem—a phenotype that is suppressed by mutation of *ZmWUS1*, indicating that REL2 and RELK act as negative regulators of meristem maintenance by modulating *ZmWUS1* activity (Liu et al. 2019; Gregory et al. 2024). Multiple *rel2*/*relk* mutants lack the embryonic SAM, demonstrating that these genes are also required for embryonic SAM establishment and maintenance (Gregory et al. 2024). Furthermore, *rel2* fails to produce axillary buds at the upper nodes of the main stem, suggesting that *REL2* is involved in the formation of axillary meristems (Liu et al. 2019).

The rice ortholog of *REL2*, *ABERRANT SPIKELET AND PANICLE1* (*ASP1*), was originally identified as a gene whose loss-of-function mutant exhibits floral and inflorescence abnormalities (Yoshida et al. 2012). *asp1* was later isolated as an enhancer of the *fon2* mutant, which exhibits an enlarged flower meristem due to stem cell overproliferation (Suzuki et al. 2019). Whereas *asp1* shows no major change in meristem size, the *fon2 asp1* double mutant displays greater enlargement of the SAM, inflorescence meristem, and flower meristem compared with *fon2* alone. Transcriptome analysis has revealed that ASP1 regulates stem cell proliferation by repressing the expression of gene sets controlled by FON2. In addition to reproductive meristems, *ASP1* is expressed during axillary meristem formation (Yoshida et al. 2012; Suzuki et al. 2019), although its role in this process remains unclear.

In this study, to better understand the mechanisms underlying stem cell proliferation during axillary meristem formation in rice, we have conducted molecular genetic analyses, focusing on the downstream mechanisms regulated by TAB1 and ASP1. Our results suggest that ASP1 acts as a novel regulator of axillary meristem formation, and that TAB1 and ASP1 antagonistically regulate cytokinin signaling to maintain proper stem cell numbers during this process.

## Results

### The stem cell region is expanded during axillary meristem formation in *asp1*

To address whether *ASP1* is involved in axillary meristem formation, we analyzed the phenotype of the *asp1-10*, a previously described loss-of-function mutant with a single nucleotide substitution in the donor splice site of intron 14 in *ASP1* (Suzuki et al 2019). First, we measured the size of the pre-meristem zone and axillary meristem of 1-month-old plants, finding that both were significantly larger in *asp1-10* than in wild type (Fig. 1, A–F). Next, we examined the spatial expression pattern of the stem cell marker *FON2* (Suzaki et al. 2006; Tanaka and Hirano 2020). In wild type, *FON2* expression was restricted to the upper domain of the pre-meristem zone (Fig. 1G), consistent with a previous report (Tanaka and Hirano 2020). In *asp1-10*, by contrast, *FON2* expression was expanded across a broader region of the pre-meristem zone (Fig. 1H), indicating expansion of the stem cell region. Quantitative real-time PCR (qRT-PCR) analysis showed that expression of *FON2* was approximately twofold higher in *asp1-10* than in wild type (Fig. 1I). Collectively, these results suggest that *ASP1* negatively regulates stem cell proliferation during axillary meristem formation.

**Figure 1.**
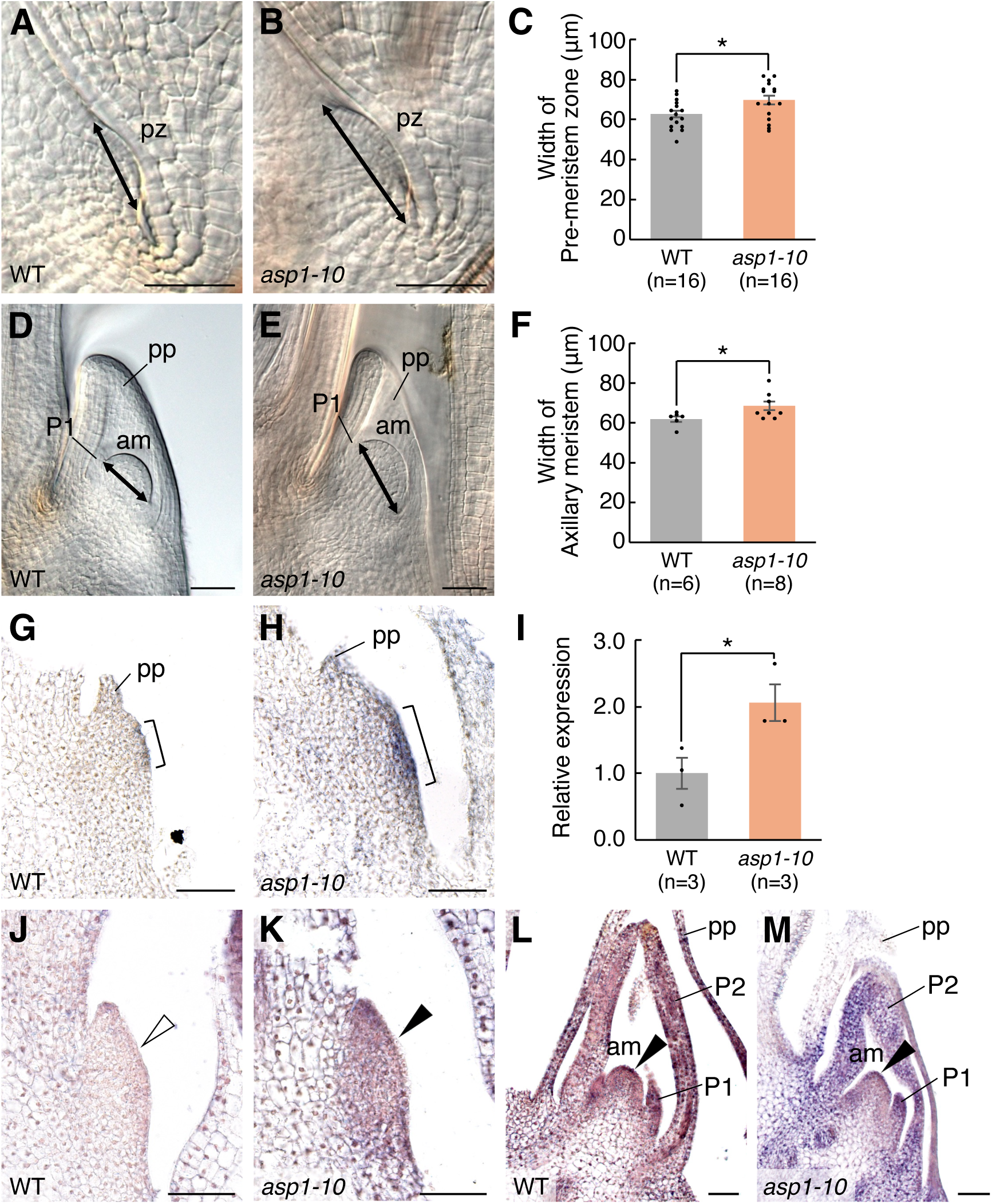
Effect of the *asp1-10* mutation on stem cell proliferation during axillary meristem formation. **A, B)** Phenotypes of the pre-meristem zone in wild type (WT) and *asp1-10*. Double-headed arrows indicate the width of the pre-meristem zone. **C)** Comparison of the width of the pre-meristem zone between WT and *asp1-10*. **D, E)** Phenotypes of the axillary meristem in WT and *asp1-10*. Double-headed arrows indicate the width of the axillary meristem. **F)** Comparison of the width of the axillary meristem between WT and *asp1-10*. **G, H)** Expression patterns of *FON2* in the pre-meristem zone in WT and *asp1-10*. Brackets indicate the expression domains of *FON2*. **I)** Relative expression level of *FON2* between wild type and *asp1-10*. **J, K)** Expression patterns of *WOX4* in the pre-meristem zone in WT and *asp1-10*. Open arrowhead indicates the absence of *WOX4* expression in the pre-meristem zone in WT; filled arrowhead indicates *WOX4* expression in *asp1-10*. **L, M)** Expression patterns of *WOX4* in the axillary meristem in WT and *asp1-10*. Filled arrowheads indicate the expression domains of *WOX4*. am, axillary meristem; pp, prophyll; pz, pre-meristem zone; P1 and P2, plastochron one-leaf and two-leaf primordium, respectively. Scale bars, 50 μm. Asterisk indicates a significant difference (P < 0.05, Student’s t-test). Error bars indicate standard error. **Alt text:** Images and bar graphs show the sizes of the pre-meristem zone and the axillary meristem in wild type and *asp1*. Images display the expression pattern of stem cell markers (*FON2*, *WOX4*) in the pre-meristem zone; bar graphs quantify the expression levels. In *asp1-10*, the stem cell region is expanded during axillary meristem formation, resulting in a larger meristem.

To examine why the stem cell region is expanded in *asp1-10*, we analyzed the expression of two stem cell-promoting factors, *TAB1* and *WOX4*, during axillary meristem formation (Ohmori et al. 2013; Tanaka et al. 2015). No significant difference in *TAB1* expression was detected between wild type and *asp1-10* (Supplemental Fig. S2). In contrast, *WOX4* expression was detected precociously in the pre-meristem zone in *asp1-10*, but was absent from this region in wild type (Fig. 1J and K). In the axillary meristem of *asp1-10*, however, *WOX4* expression was detected similarly to that in wild type (Fig. 1L and M). These observations suggest that the expanded stem cell region in *asp1-10* may be associated with the precocious expression of *WOX4*.

### *asp1* mutation rescues the defect of axillary meristem formation in *tab1*

A previous study showed that *asp1* mutation alleviates the low tiller number in *dc1*, a weak allele of *tab1* that forms a reduced number of tillers (Xia et al. 2020), because the impaired axillary bud outgrowth in *dc1* is restored by *asp1* mutation. However, it is unknown whether *asp1* mutation would affect the defect in axillary meristem formation in *tab1*. To address this issue, we generated a *tab1-1 asp1-10* double mutant and analyzed its phenotype. Six weeks after germination, *asp1-10* formed a similar number of tillers as wild type (Fig. 2A, B, and E), whereas *tab1-1* formed no tillers (Fig. 2C and E), as reported previously (Tanaka et al. 2015). In contrast to *tab1-1*, the *tab1-1 asp1-10* double mutant formed a few tillers (Fig. 2D and E), indicating that the *asp1-10* mutation partially rescued the tiller defects in *tab1-1*.

**Figure 2.**
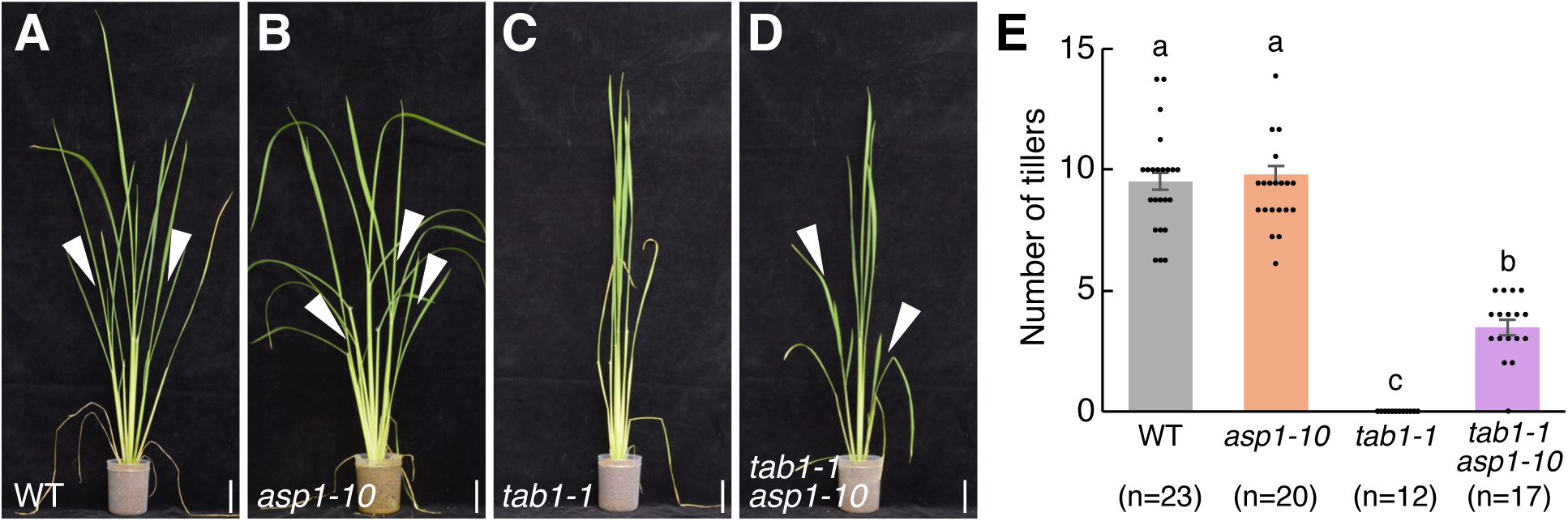
Comparison of shoot phenotypes. **A–D)** Shoot phenotypes of wild type (WT), *asp1-10*, *tab1-1*, and *tab1-1 asp1-10*, 6 weeks after germination. Arrowheads indicate tillers. **E)** Number of tillers in WT, *asp1-10*, *tab1-1*, and *tab1-1 asp1-10*. Scale bars, 5 cm. The letters a, b, and c indicate significant differences between genotypes (P < 0.01, Tukey’s test). Error bars indicate standard error. **Alt text:** Images and bar graphs show the number of tillers in wild type, *tab1-1*, *asp1-10*, and *tab1-1 asp1-10*. *asp1-10* forms a similar number of tillers as wild type. Whereas *tab1-1* has no tillers, the *tab1-1 asp1-10* double mutant forms tillers.

Next, we observed the morphology of the prophyll, which encloses the axillary bud, and its inner structure. Whereas the prophyll of *asp1-10* was similar to that of wild type (Fig. 3A and B), loss or an abnormally elongated prophyll was observed in *tab1-1* (Fig. 3C), as described previously (Tanaka et al. 2015). Notably, a wild-type-like prophyll was formed in *tab1-1 asp1-10* (Fig. 3D). Inside the prophyll of *tab1-1 asp1-10*, a dome-shaped axillary meristem, similar to that in wild type, was observed together with leaf primordia in many plants (Fig. 3E, F, H and Q); however, no meristem was detected in *tab1-1* (Fig. 3G and Q). Consistent with this, the meristem marker *OSH1* was clearly expressed during axillary meristem formation in *tab1-1 asp1-10*, similar to its expression in wild type (Fig. 3I–P). Collectively, these results indicate that the failure of axillary meristem formation in *tab1-1* is rescued by the *asp1-10* mutation.

**Figure 3.**
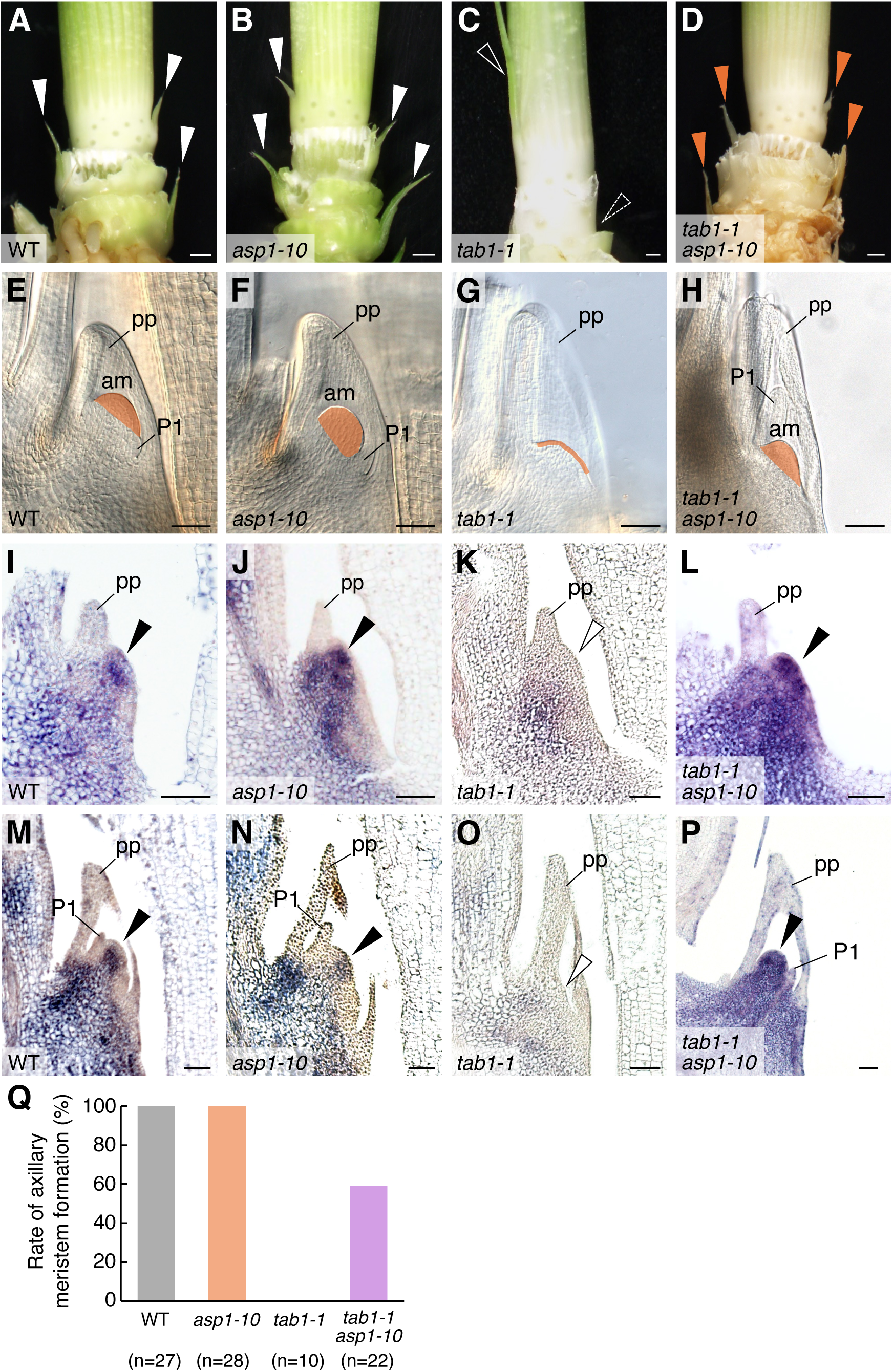
Comparison of axillary meristem phenotypes. **A–D)** Basal regions of the main stems of wild type (WT), *asp1-10*, *tab1-1*, and *tab1-1 asp1-10*, 6 weeks after germination. White filled arrowheads indicate prophylls; dashed arrowhead indicates loss of a prophyll; white open arrowhead indicates an abnormally elongated prophyll; and orange arrowheads indicate WT-like prophylls. **E–H)** Inner structure of the prophyll in WT, *asp1-10*, *tab1-1*, and *tab1-1 asp1-10*. Orange regions indicate an axillary meristem; orange line indicates its absence. **I–P)** Spatio-temporal expression patterns of *OSH1* in the pre-meristem zone **I–L)** and axillary meristem **M–P)** in WT, *asp1-10*, *tab1-1*, and *tab1-1 asp1-10*. Filled arrowheads indicate clear *OSH1* expression; open arrowheads indicate its absence. **Q)** Comparison of the rate of axillary meristem formation among WT, *asp1-10*, *tab1-1*, and *tab1-1 asp1-10*. am, axillary meristem; pp, prophyll; P1, plastochron one-leaf primordium. Scale bars, 1 mm **A–D)** and 50 µm **E–P)**. **Alt text:** Images show the outer and inner structures of axillary buds in wild type, *tab1-1*, *asp1-10*, and *tab1-1 asp1-10*. Additional images display the expression pattern of a meristem marker in each genotype. Bar graph shows the frequency of axillary meristem formation. Unlike *tab1-1*, the *tab1-1 asp1-10* double mutant frequently forms axillary meristems.

### Transcriptome analysis of genes regulated by *TAB1* and *ASP1*

To gain insight into the downstream mechanisms regulated by TAB1 and ASP1, we analyzed the genome-wide expression profiles of wild type, *tab1-1*, *asp1-10*, and *tab1-1 asp1-10* by RNA sequencing (RNA-seq) of RNA extracted from tissues containing the pre-meristem zone collected from the basal shoots of 6-week-old plants (Supplemental Fig. S3). We then analyzed the RNA-Seq data with BGI’s Dr. Tom software to identify differentially expressed genes (DEGs) between the wild type and each mutant genotype (*tab1-1*, *asp1-10*, and *tab1-1 asp1-10*) (false discovery rate < 0.05; Supplementary Table S1). Volcano plots of the upregulated and downregulated DEGs showed that the number of DEGs in *tab1-1* was considerably reduced by the *asp1-10* mutation (Fig. 4A–C), an observation that may be associated with the phenotypic rescue of *tab1-1* by the *asp1-10* mutation.

**Figure 4.**
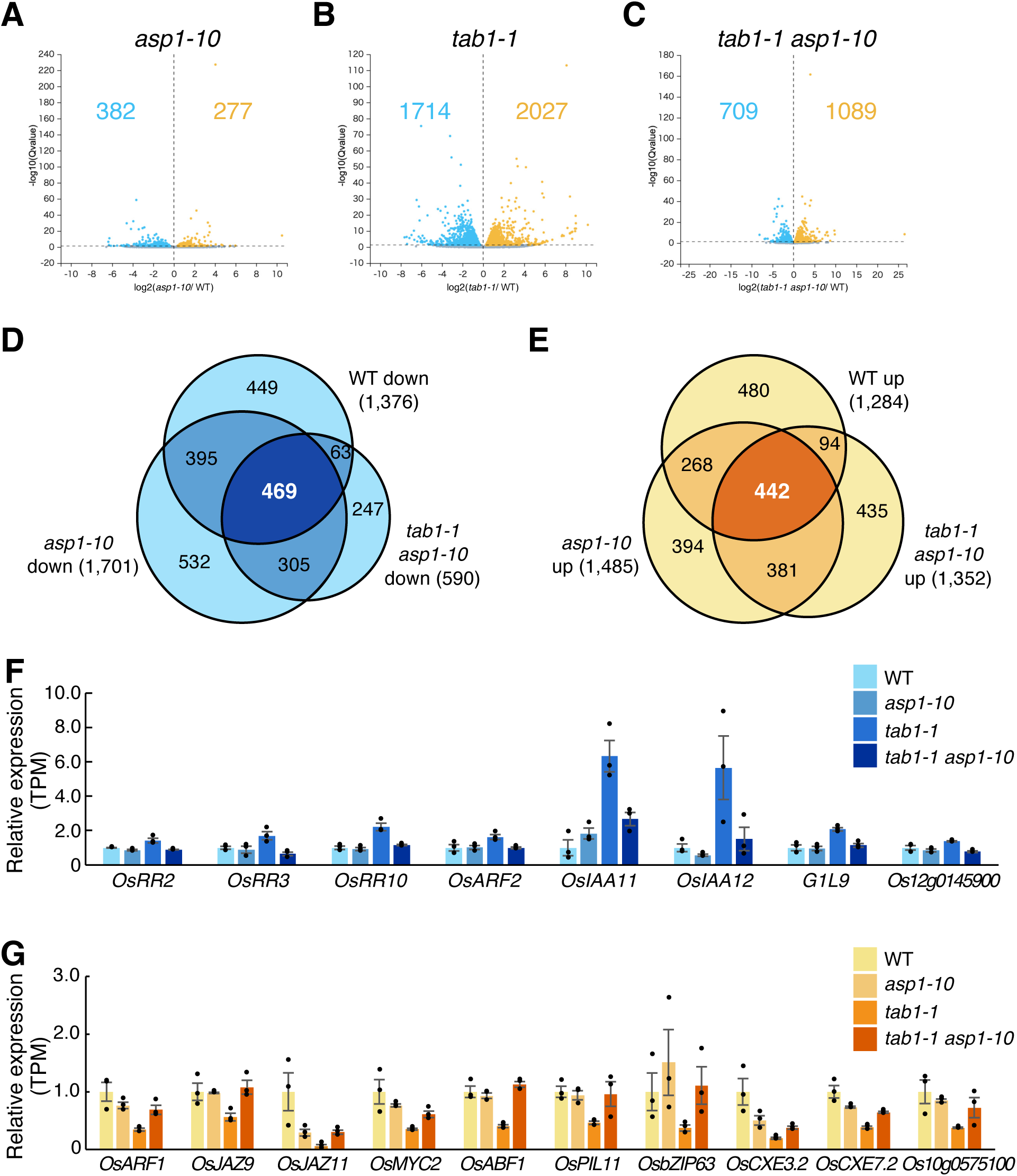
Transcriptome analysis. **A–C)** Volcano plot representation of RNA-seq analysis of *asp1-10*, *tab1-1*, and *tab1-1 asp1-10* in comparison to wild type (WT). Light blue and orange dots indicate down- and upregulated DEGs, respectively; the total numbers are also shown. Grey dots indicate non-DEGs. The screening criterion for DEGs was false discovery rate (FDR) < 0.05. **D, E)** Venn diagram of DEGs that were down- or upregulated in WT, *asp1-10,* and *tab1-1 asp1-10* relative to *tab1-1*. The screening criteria for DEGs were FDR < 0.05 and FC ≥ 1.5. **F, G)** Relative expression levels (transcripts per million; TPM) of DEGs associated with the pathway ‘plant hormone signal transduction’ that were commonly down- or upregulated in WT, *asp1-10*, and *tab1-1 asp1-10* relative to *tab1-1*. Error bars indicate standard error. **Alt text:** Volcano plots show differentially expressed genes (DEGs) in *tab1-1*, *asp1-10*, and *tab1-1 asp1-10* relative to wild type. Venn diagrams indicate shared and unique down- and upregulated DEGs relative to *tab1*. Bar graphs show the relative expression levels of DEGs associated with the pathway ‘plant hormone signal transduction’. Several type-A *OsRR* genes are upregulated in *tab1-1* but not in the other genotypes.

To identify genes involved in axillary meristem formation, we further analyzed the DEGs in three pairwise comparisons: *tab1-1* vs wild type, *tab1-1* vs *asp1-10*, and *tab1-1* vs *tab1-1 asp1-10* (false discovery rate < 0.05, fold change ≥ 1.5; Supplementary Table S2). These comparisons revealed that 469 DEGs were commonly downregulated and 442 DEGs were commonly upregulated in wild type, *asp1-10*, and *tab1-1 asp1-10* as compared with *tab1-1* (Fig. 4D and E; Supplementary Table S3). Classification of the common DEGs according to pathway terms in the Kyoto Encyclopedia of Genes and Genomes (KEGG) showed that 54 and 91 pathways were associated with the commonly downregulated and upregulated DEGs, respectively (Supplementary Table S4). Notably, among the top 15 pathways with the highest numbers of DEGs (Supplemental Fig. S4), the pathway ‘plant hormone signal transduction’ was associated with both downregulated and upregulated DEGs.

Because several plant hormones are involved in meristem function (Shi and Vernoux 2022; Hong and Fletcher 2023), we normalized and compared the expression levels of genes within this pathway among wild type, *tab1-1*, *asp1-10*, and *tab1-1 asp1-10*. Three cytokinin signaling-related genes—namely, rice type-A *RESPONSE REGULATOR2* (*OsRR2*) *OsRR3*, and *OsRR10—*as well as auxin signaling-related genes such as *OsARF2* and *IAA11* were commonly downregulated in wild type, *asp1-10*, and *tab1-1 asp1-10* as compared with *tab1-1* (Fig. 4F). By contrast, several jasmonic acid signaling-related genes such as *OsJAZ9* and *OsMYC2* were commonly upregulated in wild type, *asp1-10*, and *tab1-1 asp1-10* relative to *tab1-1* (Fig. 4G). Thus, these plant hormones may play a role in axillary meristem formation downstream of *TAB1* and *ASP1*.

Next, we normalized and compared the expression levels of genes previously implicated in axillary meristem formation, as well as rice homologs of Arabidopsis genes involved in this process, among wild type, *tab1-1*, *asp1-10*, and *tab1-1 asp1-10*. Whereas some genes (e.g., *MOC1*, *OsNAM*, *OsCUC3*) were downregulated in *tab1-1* as compared with wild type (Supplemental Fig. S5A and B), their expression did not significantly differ between *tab1-1* and *tab1-1 asp1-10*. Thus, these genes are unlikely to be related to the phenotypic rescue of *tab1-1* by the *asp1-10* mutation.

We also normalized and compared the expression levels of *ASP1*, *ASP1-RELATED1* (*ASPR1*), and *ASP2* between wild type and *tab1-1*, but found no significant difference (Supplemental Fig. S5C), indicating that *tab1* mutation does not affect the expression of these three genes.

### Ectopic expression of type-A *OsRR* genes in the pre-meristem zone in *tab1-1* but not *tab1-1 asp1-10*

As demonstrated above, KEGG pathway term classification revealed that three type-A *OsRR* genes (*OsRR2*, *OsRR3*, and *OsRR10*), which act in cytokinin signaling, were significantly upregulated in *tab1-1* but were restored to wild-type expression levels in *tab1-1 asp1-10* (Fig. 4F). Cytokinin signaling is essential for stem cell proliferation in plant meristems (Wybouw and Rybel 2019; Hong and Fletcher 2023). Notably, *WUS*, the Arabidopsis ortholog of *TAB1*, positively regulates cytokinin signaling by directly repressing the transcription of type-A ARR genes (Leibfried et al. 2005, Zhao et al. 2010). We therefore hypothesized that the three type-A *OsRR* genes may function downstream of TAB1 and ASP1.

To confirm the results of the transcriptome analysis, we examined the spatial expression patterns of *OsRR3* and *OsRR10* by *in situ* hybridization. In wild type, *asp1-10*, and *tab1-1 asp1-10*, the expression of both genes was nearly undetectable in the pre-meristem zone (Fig. 5A, B, D, E, F, and H). In contrast, clear expression of both genes was detected in the pre-meristem zone in *tab1-1* (Fig. 5C and G). Given that type-A *OsRRs* are known to negatively regulate cytokinin signaling (Leibfried et al. 2005, Zhao et al. 2010), this result suggests that cytokinin signaling may be repressed by *OsRRs* in the pre-meristem zone in *tab1-1*, but not in the other three genotypes.

**Figure 5.**
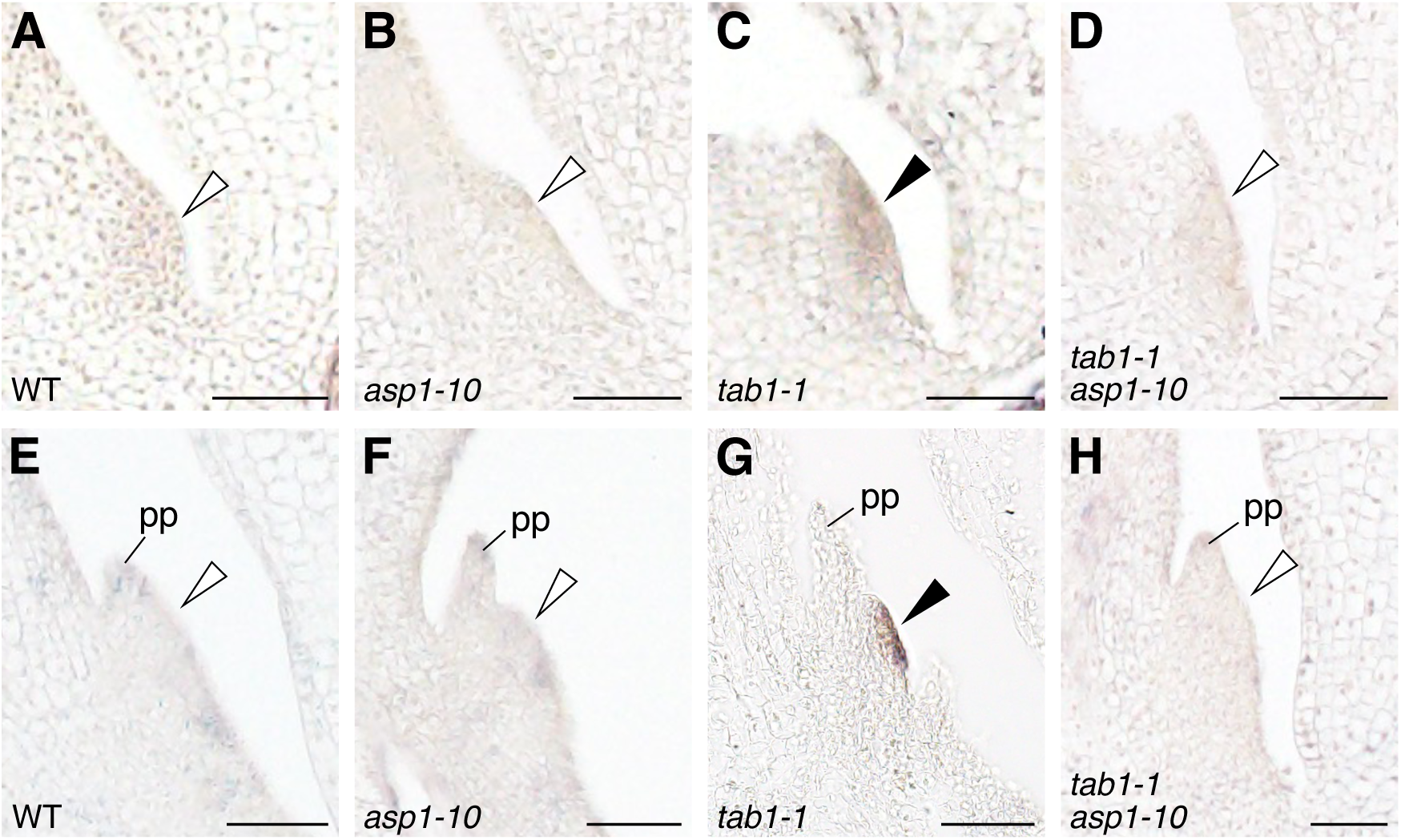
Spatio-temporal expression patterns of type-A *OsRR* genes in the pre-meristem zone. **A–D)** Expression patterns of *OsRR10* in the pre-meristem zone in wild type (WT), *asp1-10*, *tab1-1*, and *tab1-1 asp1-10*. **E–H)** Expression patterns of *OsRR3* in the pre-meristem zone in WT, *asp1-10*, *tab1-1*, and *tab1-1 asp1-10*. Open arrowhead indicates nearly undetectable expression; filled arrowhead indicates clear expression. pp, prophyll. Scale bars, 50 μm. **Alt text:** Images show the expression pattern of type-A *OsRR* genes in the pre-meristem zone of wild type, *tab1-1*, *asp1-10*, and *tab1-1 asp1-10*. Clear expression is detected in the pre-meristem zone in *tab1-1*, but not in the other genotypes.

### Cytokinin signaling is inactivated in *tab1* but restored by *asp1* mutation

To monitor cytokinin signaling outputs in plants, we introduced the synthetic cytokinin reporter *TCSv2:Clover* into wild type, *tab1-1*, and *tab1-1 asp1-1*, and selected transgenic lines in which Clover fluorescence was clearly detected in the SAM (Fig. 6A, D, and G), as reported previously (Sato et al. 2025). During axillary meristem formation in wild type, Clover fluorescence was initially observed in the pre-meristem zone (Fig. 6B). As the dome-shaped axillary meristem became visible, fluorescence was observed in the upper region of the meristem (Fig. 6C), suggesting that cytokinin signaling is activated during axillary meristem formation in rice. In *tab1-1*, by contrast, no Clover fluorescence was detected throughout axillary meristem formation (Fig. 6E and F), suggesting that cytokinin signaling is probably inactivated. In *tab1-1 asp1-10*, clear Clover fluorescence, similar to that in wild type, was observed in both the pre-meristem zone and the axillary meristem (Fig. 6H and I). Collectively, these observations suggest that the defect in cytokinin signaling in *tab1-1* was restored by the *asp1-10* mutation.

**Figure 6.**
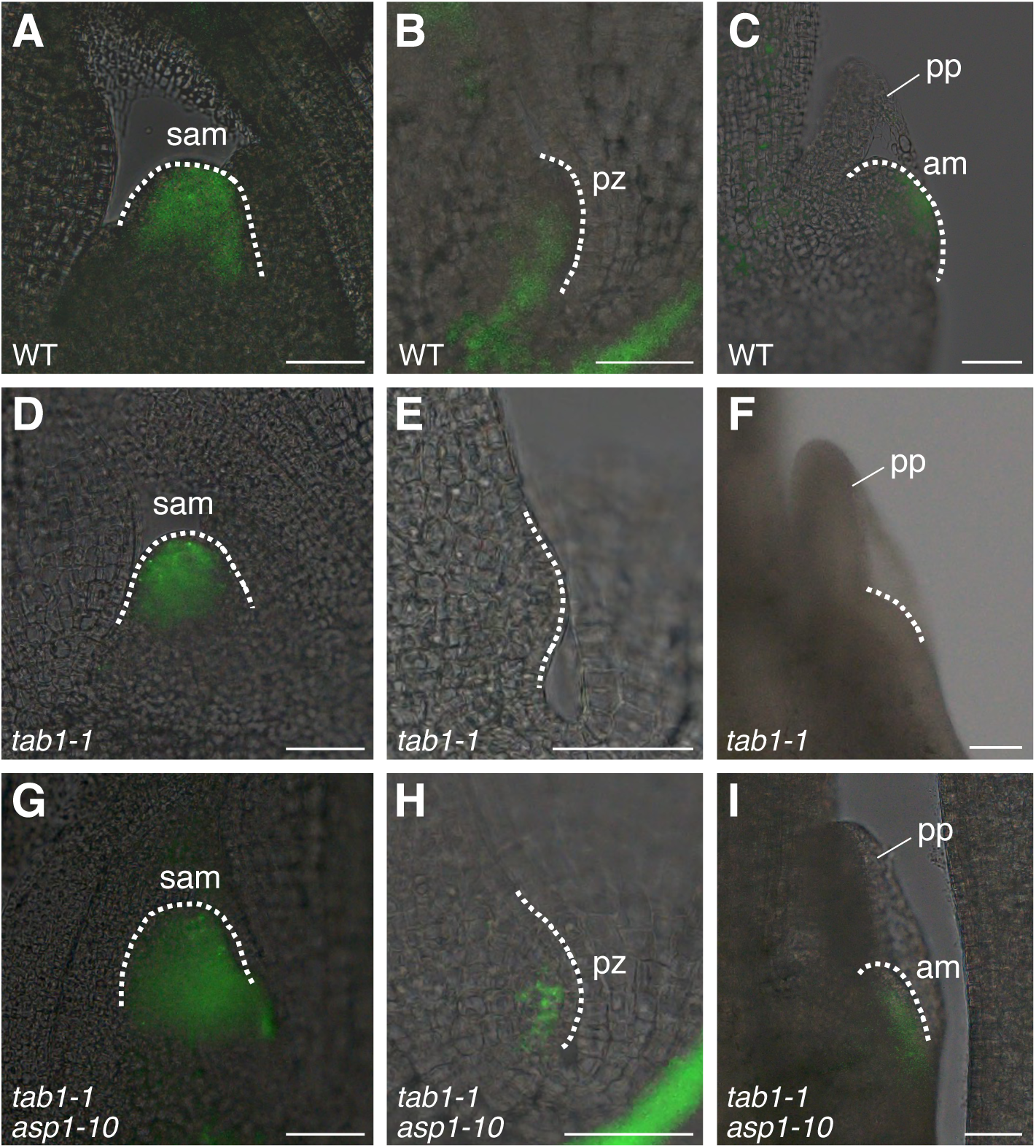
Fluorescence of TCSv2:Clover in the SAM and during axillary meristem formation. **A–C)** Fluorescence of *TCSv2:Clover* in the SAM **A)**, pre-meristem zone **B)**, and axillary meristem **C)** in wild type. **D–F)** Fluorescence of *TCSv2:Clover* in the SAM **D)**, and the region where the axillary meristem should form **E, F)** in *tab1-1*. **G–I)** Fluorescence of *TCSv2:Clover* in the SAM **G)**, pre-meristem zone **H)**, and axillary meristem **I)** in *tab1-1 asp1-10*. am, axillary meristem; pp, prophyll; pz, pre-meristem zone; sam, shoot apical meristem. Scale bars, 50 μm. **Alt text:** Images show fluorescence of the synthetic cytokinin reporter in the SAM, pre-meristem zone, and axillary meristem of wild type, *tab1-1* and *tab1-1 asp1-10*. Fluorescence is detected during axillary meristem formation in wild type and the *tab1-1 asp1-10* double mutant, but not in *tab1-1*.

### Overexpression of *OsRR10* inhibits axillary meristem formation

To gain further insight into the function of *OsRRs* and cytokinin signaling in axillary meristem formation, we generated transgenic plants overexpressing *OsRR10* during axillary meristem formation under the control of the rice *FINE CULM1* (*FC1*) promoter in both wild-type and *tab1-1 asp1-10* backgrounds (*pFC1*-*OsRR10*/WT and *FC1*-*OsRR10*/*tab1-1 asp1-10*) (Minakuchi et al. 2010; Tanaka et al. 2024). We obtained two independent lines each for *pFC1*-*OsRR10*/WT and *FC1*-*OsRR10*/*tab1-1 asp1-10* and compared their shoot phenotypes with those of wild type, *tab1-1*, and *tab1-1 asp1-10*.

Whereas wild-type control plants formed a few tillers, *pFC1*-*OsRR10*/WT plants failed to form tillers (Fig. 7A–D). Furthermore, abnormal morphology was frequently observed in the prophyll of *pFC1*-*OsRR10*/WT plants. Approximately one-third of the plants showed a reduction in meristem size, while another third completely lacked a meristem, resembling the phenotype observed in *tab1-1* (Fig. 7E–H). These observations suggest that stem cell proliferation during axillary meristem formation is probably compromised by the overexpression of *OsRR10*.

**Figure 7.**
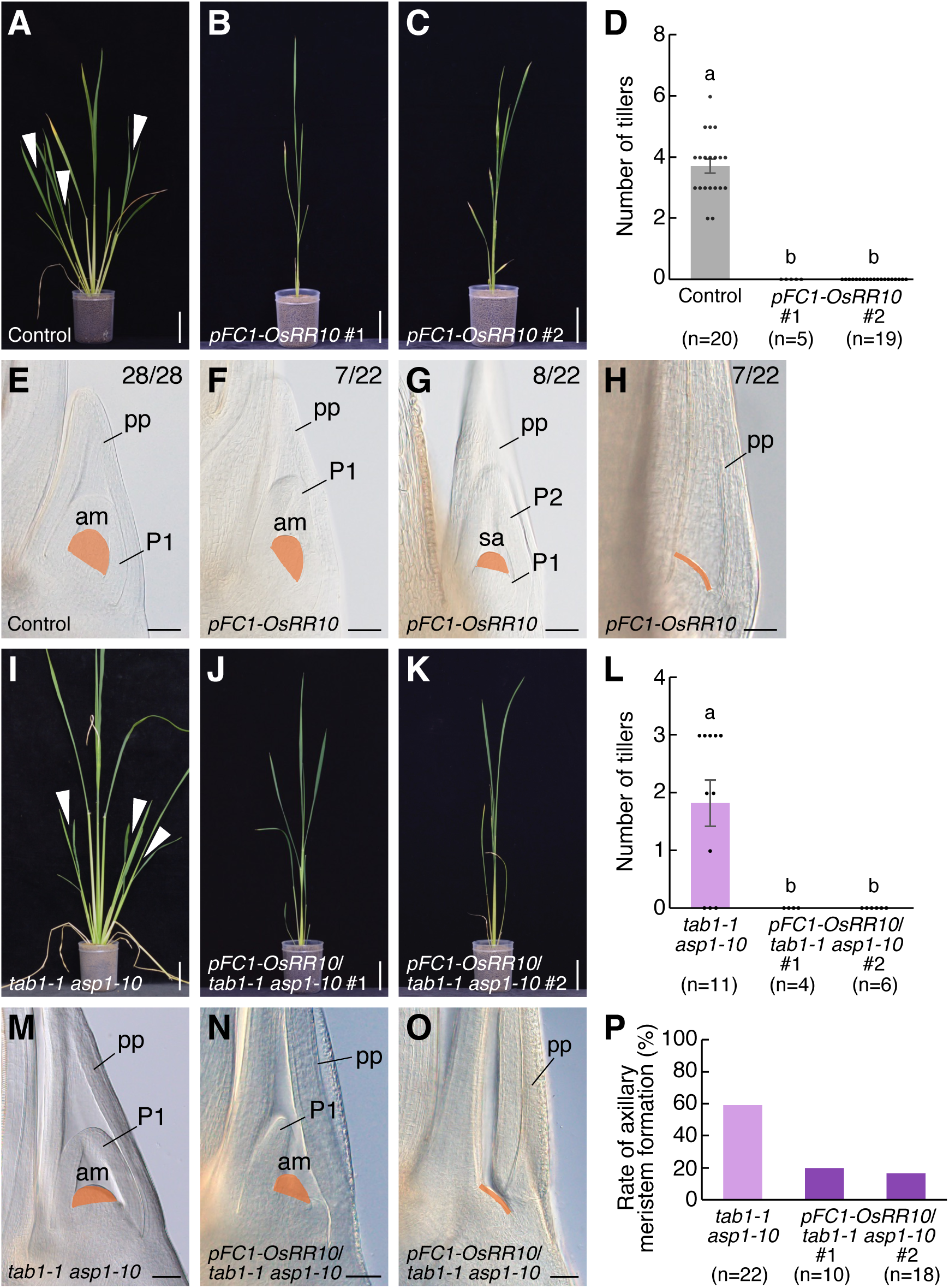
Effect of *OsRR10* overexpression on tiller and axillary meristem formation. **A–C)** Shoot phenotypes of control and *pFC-OsRR10/*WT plants, ∼7 weeks after regeneration. Arrowheads indicate tillers. **D)** Comparison of tiller number between control and *pFC-OsRR10*/WT plants. The letters a and b indicate significant differences between control and *pFC-OsRR10*/WT plants (P < 0.01, Tukey’s test). **E–H)** Inner structure of the prophyll in control and *pFC-OsRR10*/WT plants. Orange regions indicate an axillary meristem; orange line indicates its absence. Numbers (upper right) indicate the fraction of plants showing each phenotype among total prophylls examined. **I–K)** Shoot phenotypes of *tab1-1 asp1-10* and *pFC-OsRR10*/*tab1-1 asp1-10* plants, ∼7 weeks after regeneration. Arrowheads indicate tillers. **L)** Comparison of tiller number in *tab1-1 asp1-10* and *pFC-OsRR10*/*tab1-1 asp1-10* plants. The letters a and b indicate significant differences between *tab1-1 asp1-10* and *pFC-OsRR10*/*tab1-1 asp1-10* plants (P < 0.05, Tukey’s test). **M–O)** Inner structure of the prophyll in *tab1-1 asp1-10* and *pFC-OsRR10*/*tab1-1 asp1-10* plants. Orange regions indicate an axillary meristem; orange line indicates its absence. **P)** Comparison of the rate of axillary meristem formation between *tab1-1 asp1-10* and *pFC-OsRR10*/*tab1-1 asp1-10* plants. am, axillary meristem; pp, prophyll; P1 and P2, plastochron one-leaf and two-leaf primordium, respectively; sa, small axillary meristem. Scale bars, 5 cm **A–C, I–K)** and 50 µm **E–H, M–O)**. Error bars indicate standard error. **Alt text:** Images show shoots and inner structures of axillary buds, and bar graphs show the number of tillers in *pFC-OsRR10*/WT and *pFC-OsRR10*/*tab1-1 asp1-10* plants. A bar graph also shows the frequency of axillary meristem formation in *pFC-OsRR10*/*tab1-1 asp1-10* plants. Overexpression of *OsRR10* inhibits axillary meristem formation in wild type and considerably reduces its frequency in the *tab1-1 asp1-10* double mutant.

In contrast to *tab1-1 asp1-10* plants, *pFC1*-*OsRR10*/*tab1-1 asp1-10* plants failed to form tillers (Fig. 7I–L) and frequently lacked axillary meristems (Fig. 7M–P), indicating that *asp1* mutation does not rescue the *tab1* phenotype when *OsRR10* is overexpressed.

## Discussion

Little was previously known about how *TAB1* promotes stem cell proliferation during axillary meristem formation in rice, but this study has demonstrated that *TAB1* fulfills this role by activating cytokinin signaling. In addition, we have identified *ASP1* as a novel factor that negatively regulates stem cell proliferation during axillary meristem formation. *ASP1* probably restricts stem cell proliferation by repressing the expression of *WOX4*, a paralog of *TAB1*. Furthermore, transcriptome and genetic analyses using a *tab1 asp1* double mutant revealed that *ASP1* negatively regulates cytokinin signaling by promoting the expression of type-A *OsRR* genes.

### *TAB1* promotes stem cell proliferation by activating cytokinin signaling

Cytokinins play a critical role in regulating stem cell homeostasis. In Arabidopsis, cytokinin signaling promotes stem cell proliferation in meristems in association with the function of WUS (Wybouw and Rybel 2019; Hong and Fletcher 2023). During axillary meristem formation, cytokinin signaling—as monitored by the TCS reporter—is detected in the leaf axil prior to any morphological changes, and subsequently induces *WUS* expression as the meristem is established (Wang et al. 2014; Wang et al. 2017). In turn, WUS positively regulates cytokinin signaling (Leibfried et al. 2005; Zhao et al. 2010), resulting in a positive feedback loop between cytokinin signaling and WUS. The axillary meristem is frequently compromised in mutants of the cytokinin receptor *AHK* or the type-B *ARRs* (Wang et al. 2014), indicating the importance of cytokinin signaling in its formation. Despite these findings in Arabidopsis, the role of cytokinin signaling in axillary meristem formation in rice remains unclear.

In this study, we utilized the TCSv2:Clover reporter to monitor cytokinin signaling outputs during axillary meristem formation in rice. In the wild-type background, Clover fluorescence was detected initially in the pre-meristem zone and later in the fully developed axillary meristem. As previously reported, the pre-meristem zone is the site where stem cells are established during axillary meristem formation (Tanaka and Hirano 2020). We observed that Clover fluorescence was largely confined to this zone, overlapping with the stem cell region. Once the meristem dome was fully formed, Clover fluorescence was localized to the apical region of the meristem, where stem cells are maintained (Tanaka and Hirano 2020). These findings suggest that cytokinin signaling is probably involved in stem cell proliferation during axillary meristem formation in rice.

Transcriptome analysis identified three type-A *OsRR* genes (*OsRR2*, *OsRR3*, *OsRR10*) whose expression levels were significantly upregulated in *tab1* as compared with other genotypes. In Arabidopsis, WUS activates cytokinin signaling by directly repressing the transcription of several type-A *ARR* genes (*ARR5–ARR7*, *ARR15*) (Leibfried et al. 2005; Zhao et al. 2010). Given this fact, the three type-A OsRRs were considered to act downstream of TAB1.

Spatio-temporal analysis further revealed that these *OsRR* genes were expressed in the pre-meristem zone in *tab1*, and their expression domains overlapped with that of *TAB1* observed in the wild type (Tanaka et al. 2015). In contrast, the expression of the *OsRR* genes was not detected in the pre-meristem zone in wild type, *asp1-10*, or *tab1-1 asp1-10*. Furthermore, numerous WUS/TAB1-binding motifs were found in the promoter regions of the *OsRR* genes (Supplemental Fig. S6) (Leibfried et al. 2005). These findings suggest that TAB1 probably activates cytokinin signaling by repressing the transcription of the type-A *OsRR* genes. To support this idea, we analyzed TCSv2:Clover fluorescence in *tab1* and found that fluorescence was absent during axillary meristem formation. Furthermore, when *OsRR10* overexpression was driven by the *FC1* promoter during axillary meristem formation (*pFC1-OsRR10*/WT), axillary meristem formation was frequently inhibited, partially phenocopying *tab1*. In Arabidopsis, overexpression of *ARR7*, a target of WUS, leads to termination of the SAM—a phenotype very similar to that of *wus* (Leibried et al. 2005). This is consistent with our observation that overexpression of *OsRR10* lead to a *tab1*-like phenotype. Therefore, the phenotype of *pFC1-OsRR10*/WT seems likely to result from the repression of cytokinin signaling due to *OsRR10* overexpression in the pre-meristem zone, probably leading to failure of stem cell proliferation.

Collectively, our results suggest that TAB1 activates cytokinin signaling by repressing the expression of type-A *OsRRs* in the pre-meristem zone during axillary meristem formation (Supplemental Fig. S7B). This activation of cytokinin signaling is likely to be essential for promoting stem cell proliferation and facilitating axillary meristem formation. Therefore, the role of WUS in regulating stem cell proliferation by activating cytokinin signaling in meristems in Arabidopsis seems to be conserved in TAB1 during axillary meristem formation in rice.

### *ASP1* acts as a negative regulator of stem cell proliferation during axillary meristem formation

In this study, we showed that meristem size during axillary meristem formation was significantly larger in *asp1* than in wild type. In addition, we observed expansion of the stem cell region and elevated expression of *FON2* in *asp1* as compared with wild type. These findings suggest that *ASP1* plays a critical role in restricting stem cell proliferation during axillary meristem formation.

What is the mechanism by which *ASP1* restricts stem cell proliferation? Analysis of *TAB1* mRNA during axillary meristem formation showed that expression levels in *asp1* were comparable to those in wild type. In contrast, there was a distinct difference in the spatio-temporal expression pattern of *WOX4* between wild type and *asp1*: namely, *WOX4* expression was precociously detected at an early stage of axillary meristem formation in *asp1*. In wild type, *WOX4* is expressed at a later stage of axillary meristem formation when *TAB1* expression has disappeared, and it is required for stem cell proliferation at that stage (Supplemental Fig. S1) (Ohmori et al. 2013; Tanaka et al. 2015). In *asp1*, therefore, it seems likely that the simultaneous expression of two stem cell-promoting factors, *TAB1* and *WOX4*, during the early stage of axillary meristem formation may be responsible for the expanded stem cell region observed in this mutant. Collectively, these findings suggest that *ASP1* negatively regulates stem cell proliferation, probably by repressing the precocious expression of *WOX4* during axillary meristem formation (Supplemental Fig. S7A).

We previously reported that *ASP1* negatively regulates stem cell proliferation in the SAM, inflorescence meristem, and flower meristem (Suzuki et al. 2019). Interestingly, in contrast to the axillary meristem, the size of the inflorescence meristem in the *asp1* single mutant is comparable to that in wild type. One possible explanation for this difference is that *TAB1* and *WOX4* are expressed at similar levels in the inflorescence meristem of both wild type and *asp1* (Suzuki et al. 2019). Therefore, *ASP1* is unlikely to be involved in *WOX4* repression in the inflorescence meristem, suggesting that a factor required for *ASP1*-mediated repression of *WOX4* may be absent in this meristem.

The role of *ASP1*-like genes in axillary meristem formation has been suggested in other plant species. In maize, a loss-of-function mutant of *REL2*, an *ASP1* ortholog, exhibits a partial loss of axillary buds (Liu et al. 2019), suggesting that *REL2* likely promotes axillary meristem formation. Similarly, in Arabidopsis, mutants with multiple mutations of *TPL/TPR* genes, homologs of *ASP1*, lack axillary meristems in most cases (Liu et al. 2024)—a phenotype that contrasts with rice *asp1*. *ASP1*, *REL2*, and *TPL* each encode a transcriptional corepressor of the Groucho/Tup1 family that regulates target gene expression through interactions with various transcription factors (Kieffer et al. 2006; Long et al. 2006; Liu and Karmarker 2008; Yoshida et al. 2012; Liu et al. 2019; Plant et al. 2021). Thus, the distinct axillary meristem phenotypes observed in rice *asp1*, maize *rel2*, and Arabidopsis *tpl* may reflect differences in the interacting partners or downstream target genes of ASP1, REL2, and TPL.

### *ASP1* restricts stem cell proliferation by negatively regulating cytokinin signaling

This study has revealed that *asp1* mutation rescues the defect of axillary meristem formation in *tab1*, raising questions about the underlying mechanism. Transcriptome analysis revealed that three type-A *OsRR* genes (*OsRR2*, *OsRR3*, *OsRR10*) were upregulated in *tab1*. In *tab1 asp1*, by contrast, their expression was restored to wild-type levels and was undetectable in the pre-meristem zone. Consistent with this, TCSv2:Clover fluorescence was clearly observed in the pre-meristem zone in *tab1 asp1*, indicating active cytokinin signaling, whereas fluorescence was absent in *tab1*. These findings suggest that the rescue of axillary meristem formation in *tab1 asp1* results from the restoration of cytokinin signaling. In support of this idea, the introduction of *pFC1-OsRR10* into *tab1 asp1* to repress cytokinin signaling led to the inhibition of axillary meristem formation in most plants. Thus, rescue of the *tab1* phenotype by *asp1* mutation depends on cytokinin signaling regulated by type-A *OsRRs*.

Collectively, our results suggest that ASP1 negatively regulates cytokinin signaling by promoting the expression of type-A *OsRR* genes (Supplemental Fig. S7B). To date, studies examining the relationship between TPL/ASP1 and cytokinin action have been limited to reports showing that *TPL*-related genes regulate the expression of cytokinin metabolic enzymes and cytokinin-responsive genes (He et al. 2022; Jing et al. 2023). Our study provides the first evidence supporting the involvement of ASP1 in cytokinin signaling.

How, then, does ASP1 promote the expression of type-A *OsRR* genes? Given that *ASP1* encodes a Groucho/Tup1 family transcriptional corepressor (Yoshida et al. 2012), it is unlikely that ASP1 directly regulates the transcription of *OsRR* genes. Instead, ASP1 probably acts through an unknown factor ‘X’ to promote the expression of *OsRR* genes (Supplemental Fig. S7B). One potential candidate for factor X is WOX4: as described above, *WOX4* was precociously expressed during axillary meristem formation in *asp1*. Moreover, expression of *WOX4* driven by the *TAB1* promoter (*pTAB1-WOX4*) has been shown to rescue the defect of axillary meristem formation in *tab1* (Tanaka and Hirano, 2020). Thus, WOX4 is a plausible candidate for factor X. Another possible candidate is an AUXIN RESPONSE FACTOR (ARF) because Arabidopsis ARF5 has been reported to directly repress the expression of type-A *ARR* genes (Zhao et al. 2010). However, our transcriptome analysis did not identify any *ARF* genes with higher expression in *tab1 asp1* relative to *tab1* (Supplemental Fig. S5D), suggesting that *ARF* genes are unlikely to play a major role in this context. Further studies will be required to determine whether WOX4 corresponds to factor X or to identify the factor that fulfills this role.

Notably, *asp1* mutation did not fully rescue the defect of axillary meristem formation in *tab1*, although cytokinin signaling was activated during axillary meristem formation in *tab1 asp1*, similar to wild type. This observation suggests that TAB1 also promotes stem cell proliferation through a cytokinin-independent pathway that does not involve ASP1 (Supplemental Fig. S7B). In Arabidopsis, WUS maintains SAM not only by regulating cytokinin signaling but also by controlling auxin signaling and response pathways (Ma et al. 2019). TAB1 may similarly regulate stem cell proliferation during axillary meristem formation through such mechanisms in rice. Elucidating these additional pathways will be an important direction for future studies.

### Genetic relationship between *TAB1* and *ASP1*

Our results indicate that TAB1 and ASP1 antagonistically regulate cytokinin signaling during axillary meristem formation (Supplemental Fig. S7B). Phenotypic analysis of *tab1 asp1* further suggests that *asp1* is epistatic to *tab1*. In Arabidopsis, WUS forms a complex with TPL, and this complex is required for maintaining the flower meristem (Causier et al. 2012). A physical interaction between TAB1 and ASP1 has been also reported in rice (Lu et al. 2015; Xia et al. 2020); however, the assumption that TAB1 and ASP1 function as a protein complex would make it difficult to explain their antagonistic roles on cytokinin signaling and the observed epistatic relationship. This raises questions about the exact relationship between *TAB1* and *ASP1* during axillary meristem formation in rice.

One possibility is that TAB1 may repress ASP1 function. Transcriptome analysis revealed no significant difference in *ASP1* expression between wild type and *tab1* (Supplemental Fig. S5C), suggesting that TAB1 does not regulate *ASP1* at the transcriptional level. However, posttranslational regulation cannot be excluded. An Arabidopsis a homolog of ASP1, TRP1, undergoes SUMOylation, which attenuates its transcriptional repression activity (Niu et al. 2019). As our transcriptome analysis showed that genes involved in SUMOylation are downregulated in *tab1* relative to wild type (Supplemental Fig. S5E), it is possible that TAB1 may promote ASP1 SUMOylation and thereby attenuate its activity.

Alternatively, TAB1 and ASP1 may function through independent pathways. In Arabidopsis, TPL forms a complex with BRANCHED1 (BRC1) to regulate axillary bud outgrowth (Yang et al. 2018). In rice, *FC1*, an ortholog of *BRC1*, negatively regulates stem cell proliferation during axillary meristem formation through a *TAB1*-independent pathway (Tanaka et al. 2024). Based on these findings, ASP1 may interact with FC1 to regulate stem cell proliferation independently of TAB1. In support of this idea, *WOX4* is precociously expressed during axillary meristem formation in *fc1* (Tanaka et al. 2024), as observed in *asp1*. This suggests that ASP1 and FC1 may cooperatively repress *WOX4* expression.

## Materials and methods

### Plant materials and growth conditions

*Oryza sativa* L. ssp. *japonica* cultivar Taichung 65 (T65) was used as the wild type in phenotypic and *in situ* hybridization analyses. The *tab1-1* and *asp1-10* mutants have been reported previously (Tanaka et al. 2015; Suzuki et al. 2019). Plants were grown in an NK System Biotron (model LH-350S, Nippon Medical & Chemical Instruments, Osaka, Japan) at 28 °C under 14 h/10 h light/dark conditions.

### Clearing experiment

To observe the inner structures of axillary buds or prophylls, shoot tissues (2–4 mm) were dissected and fixed in a 1:3 (v/v) solution of acetic acid and ethanol. After ∼1 day, the samples were transferred into distilled water for 2 hours, and then into an 8:3:1 (v/v) transparency solution of chloral hydrate, water, and glycerol for at least 1 day. Samples were observed under differential interference contrast optics by a BX53LED optical microscope (Olympus, Tokyo, Japan) with a DP74 color camera (Olympus).

### *In situ* hybridization

The basal parts of shoot tissues containing the pre-meristem zone and axillary meristem were dissected, and then fixed and dehydrated in accordance with the methods of Toriba and Hirano (2018). The samples were placed in xylene and then embedded in Paraplast Plus (Leica, Wetzlar, Germany).

To generate *OsRR3* and *OsRR10* probes for *in situ* hybridization, cDNA fragments including the coding sequence were amplified with appropriate primers (Supplementary Table S5) and cloned into a pCRII vector (Thermo Fisher Scientific, Waltham, USA). RNA probes were transcribed with T7 or Sp6 RNA polymerase using digoxigenin-labeled UTPs (Roche Applied Science, Mannheim, Germany). The *OSH1*, *FON2*, and *WOX4* probes were prepared as described previously (Suzaki et al. 2006, Ohmori et al. 2013, Tanaka et al. 2015). Microtome sections (10 μm) were cut from tissues embedded in Paraplast Plus and mounted on glass slides. The *in situ* hybridization experiments and immunological detection of signals were performed according to the methods of Toriba and Hirano (2018). A BX53LED optical microscope (Olympus) with a DP74 color camera (Olympus) was used for observation.

### qRT-PCR analysis

For analysis of *FON2* and *TAB1* expression, total RNA was isolated from tissues containing the pre-meristem zone using a RNeasy Plant Mini Kit (Qiagen, Hilden, Germany) according to the manufacturer’s instructions. First-strand cDNA was synthesized from 500 ng of total RNA using a PrimScript RT reagent kit (TaKaRa Bio, Shiga, Japan). qRT-PCR was performed by using a StepOn Real-Time PCR System (Thermo Fisher Scientific) with Power SYBR Green PCR Master Mix (Thermo Fisher Scientific). Three RNA samples, independently isolated from five shoots, were used for qRT-PCR analysis, and the PCR reaction was conducted twice for each sample. The *ACTIN1* gene was used as a control. The primers used are listed in Supplementary Table S5.

### RNA-seq experiment and analysis

Shoot apices (diameter 1.5 mm, height 2 mm), including the pre-meristem zone and SAM were used for RNA isolation. Samples were dissected from 4–5 individual plants, with three biological replicates per genotype (wild type, *tab1-1*, *asp1-10*, *tab1-1 asp1-10*). RNA was isolated by using the RNeasy Plant Mini Kit (Qiagen), with an RNase-Free DNase Set DNaseⅠ (Qiagen) digestion step, and treated with RNase-free DNaseⅠ. cDNA library construction and sequencing were performed by the Beijing Genomics Institute using the DNBSEQ platform using a sequencing length of 100-bp paired-end reads. The sequencing data were filtered with SOAPunke (v1.5.2). The data ranged from 43.95 to 46.42 million reads per library (average 45.84 million). Clean reads were aligned to the rice reference genome IRGSP-1.0, and an average of 96.69% of reads were mapped to the reference, using HISAT2 (v2.0.4). Bowtie2 (v2.2.5) was applied to align the clean reads to the reference coding gene set. The expression levels of each gene were calculated by RSEM (v1.2.8) and differential expression analysis was performed using DESeq2. DEGs were assigned as a fold change of ≥1.5 and adjusted false discovery rate of <0.05. The sequencing data analysis, including creation of Volcano plots and Venn diagrams, as well as the classification of KEGG terms, was performed in the BGI online system (Dr. Tom).

### Generation of *TCSv2:Clover* plants and imaging

The plasmid for expression of recombinant *TCSv2:Clover* was constructed as a binary vector for plant transformation by Gateway LR recombination between the pRIT2 backbone vector and a pENTR entry plasmid carrying a *TCSv2-Clover* reporter cassette, in which the *TCSv2* sequence drove *Clover* expression, the *OsADH1* sequence functioned as a translational enhancer, and the *OsHSP* sequence served as the transcription terminator. The resultant plasmid was introduced into *Agrobacterium tumefaciens* EHA101 and transformed into calli derived from T65, Nipponbare, *tab1-1*, and *tab1-1 asp1-10* according to the method of Hiei et al. (1994). To observe TCSv2:Clover fluorescence, shoot apices of transgenic plants (T0 generation) were embedded in 5% agar and sliced into 30-μm sections using a vibratome. Clover fluorescence was observed using a BX53LED optical microscope (Olympus) with a DP74 color camera (Olympus).

### Generation of *pFC1-OsRR10* plants

To generate *pFC1-OsRR10* plants, the full-length *OsRR10* cDNA and the 5-kb upstream region of *FC1* were each amplified by using appropriate primers (Supplementary Table S5). The full-length *OsRR10* cDNA was first cloned into a pENTR D-TOPO vector (Thermo Fisher Scientific, Waltham, USA); the pFC1 fragment was then cloned into the same vector using In-Fusion (Takara Bio). For plant transformation, the *pFC1*-*OsRR10* fragment was transferred into the binary vector pBI-Hm12-GW, which contains the Gateway rfC cassette, by an LR recombination reaction (Yoshida et al. 2009). The recombinant plasmids were introduced into *Agrobacterium tumefaciens* EHA101 and transformed into calli derived from T65 and *tab1-1 asp1-10* according to the method of Hiei et al. (1994). Phenotypic analyses were performed using the T0 generation of transgenic plants.

### Accession numbers

Sequence data from this study are available in GenBank, EMBL, and DDBJ under the following accession numbers: AB638269 (*ASP1*), AB245090 (*FON2*), AB218894 (*TAB1*), JF836159 (*WOX4*), D16507 (*OSH1*), AB249662 (*OsRR2*), AB249655 (*OsRR3*), AB249660 (*OsRR10*), AB071291 (*OsARF2*), NM_001417717 (*IAA11*), AK103459 (*OsJAZ9*), AK288082 (*OsMYC2*), JF836159 (*MOC1*), AB088343 (*FC1*), XM_015787089 (*OsNAM*). All RNA-seq data have been deposited with links to BioProject accession number PRJDB42173 in the DDBJ BioProject database.

## Acknowledgements

We thank Dr. Hiro-Yuki Hirano for providing *asp1-10* and *tab1-1 asp1-10* seeds.

## Author contributions

A.O. and W.T. designed the research; A.O., T.T., M.S., H.T., and W.T. conducted the experiments; A.O. analyzed the data; A.O. and W.T. drafted the manuscript with input from all other authors.

