## Supplementary figures for "*TAB1* and *ASP1* act antagonistically on cytokinin signaling to regulate axillary meristem formation in rice"

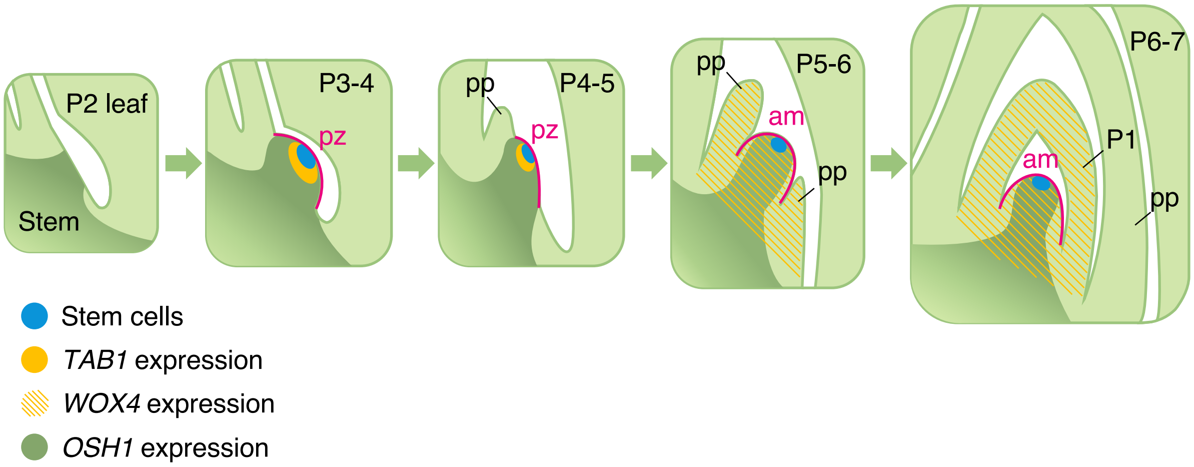


**Figure S1. Developmental stages of axillary meristem formation in rice.** Pink lines indicate the pre-meristem zone and axillary meristem. Light blue regions indicate stem cells. Yellow, yellow-hatched, and dark green regions indicate the expression domains of *TAB1*, *WOX4*, and *OSH1*, respectively. am, axillary meristem; pp, prophyll; pz, pre-meristem zone; P1 to P7, plastochron one-leaf to seven-leaf primordium, respectively.

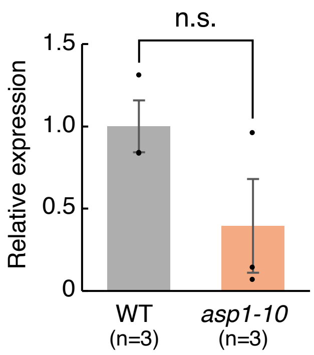


**Figure S2. Relative expression level of *TAB1* in wild type and *asp1-10*.** n.s. indicates not significant ($P \geq 0.05$, Student's t-test). Error bars indicate standard error.

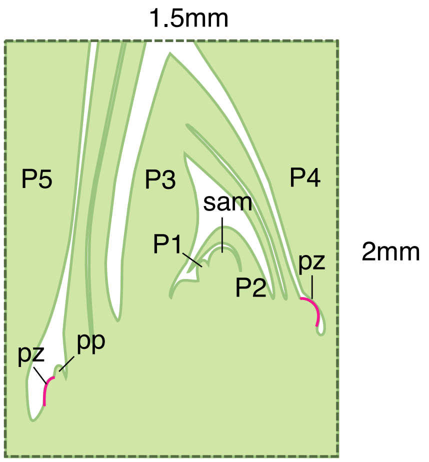


**Figure S3. Schematic of samples used in transcriptome analysis.** Pink lines indicate the pre-meristem zone. pp, prophyll; pz, pre-meristem zone; P1 to P5, plastochron one-leaf to five-leaf primordium, respectively; sam, shoot apical meristem.

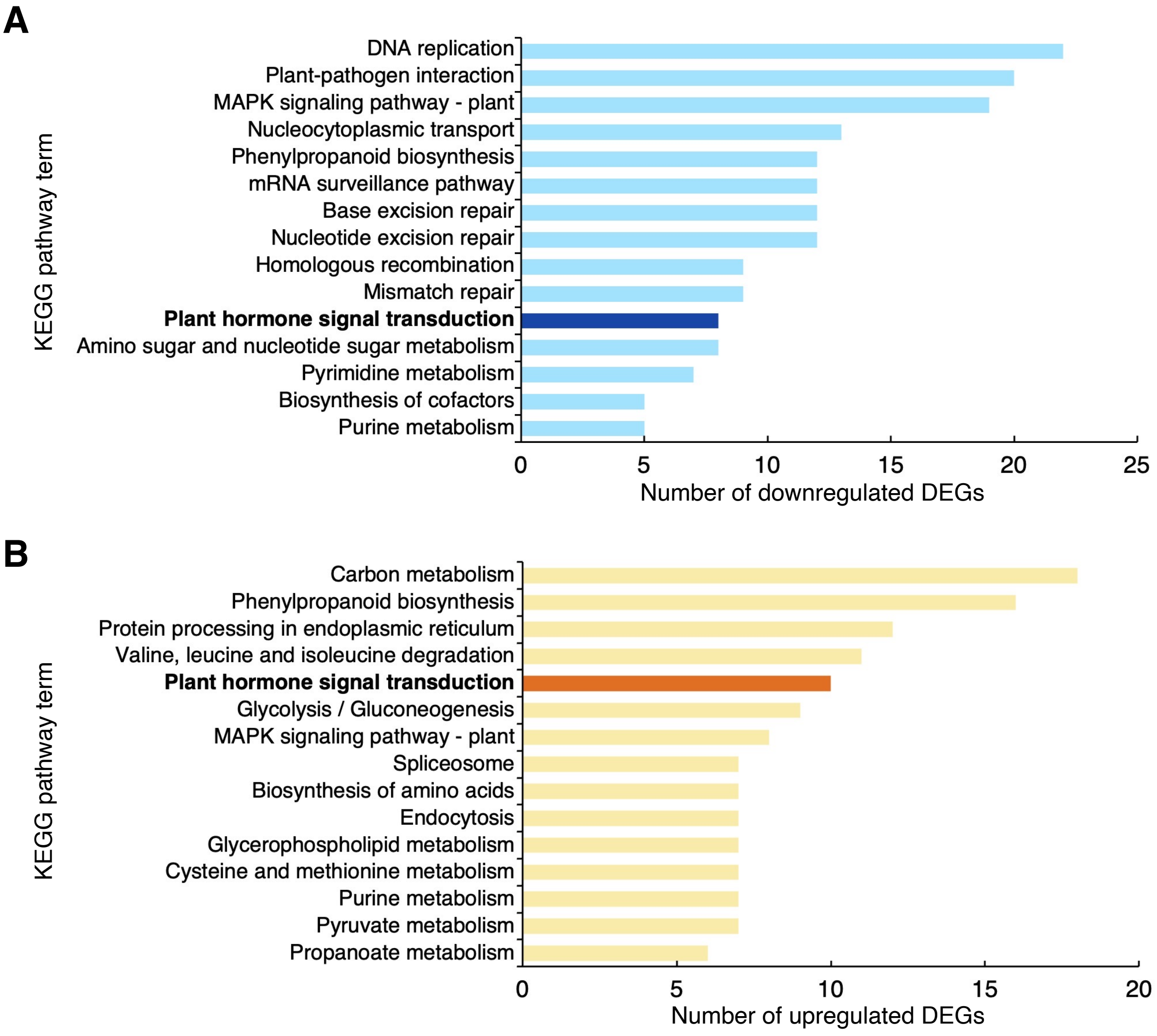


**Figure S4. KEGG pathway classification.** Top 15 KEGG pathway terms ranked by the number of commonly downregulated **A)** and upregulated **B)** DEGs in wild type, *asp1-10*, and *tab1-1 asp1-10* relative to *tab1-1*. The screening criteria for DEGs were FDR < 0.05 and FC ≥ 1.5.

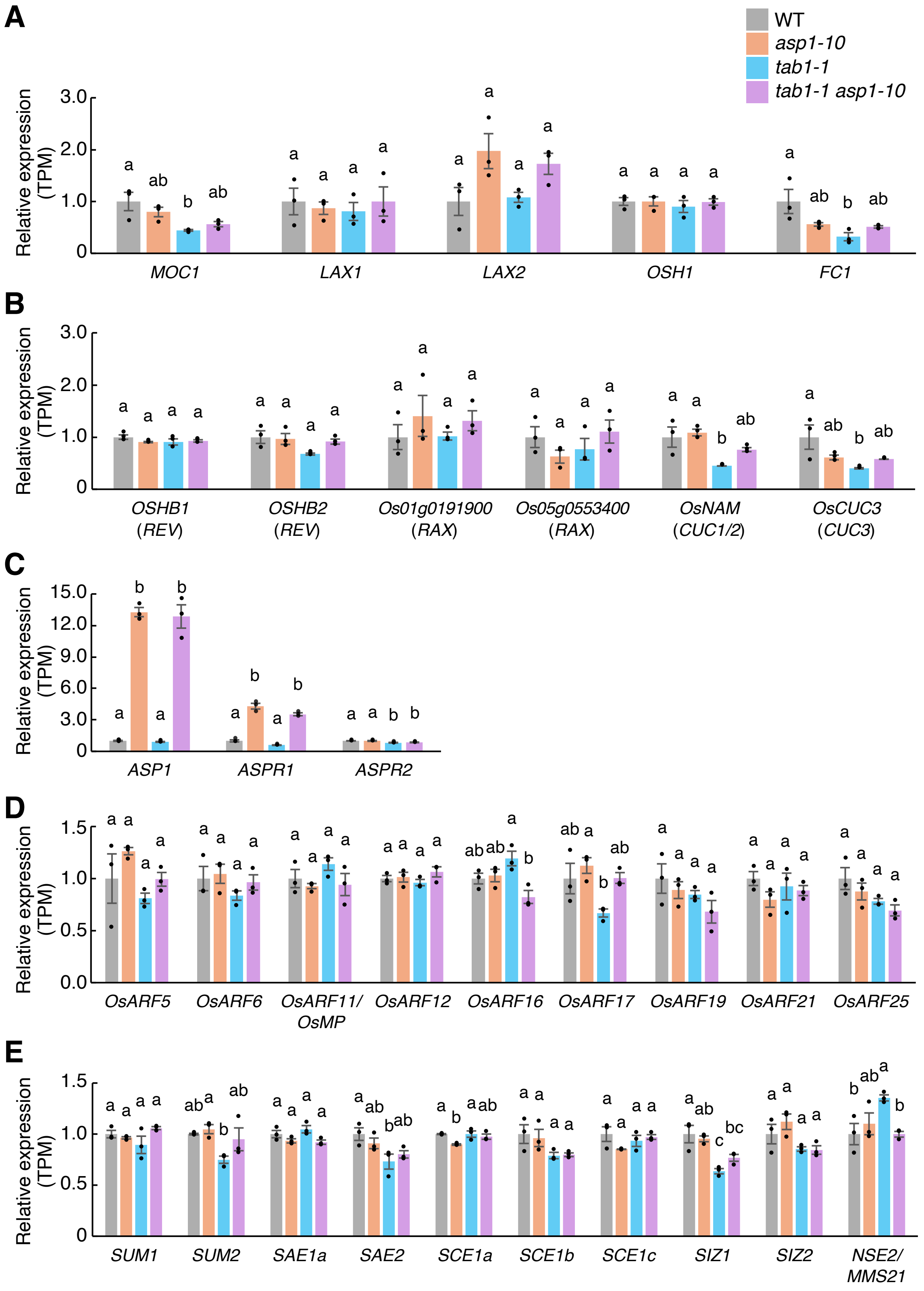


**Figure S5. Relative expression levels of genes from transcriptome data. A)** Relative expression levels (transcripts per million; TPM) of genes previously implicated in axillary meristem formation in rice. **B)** Relative expression levels of rice homologs of Arabidopsis genes involved in axillary meristem formation. **C)** Relative expression levels of *ASP1*, *ASPR1*, and *ASPR2*. **D)** Relative expression levels of ARF family genes. **E)** Relative expression levels of genes involved in SUMOylation. The letters a, b, and c indicate significant differences between genotypes ($P < 0.05$, Tukey's test). Error bars indicate standard error.

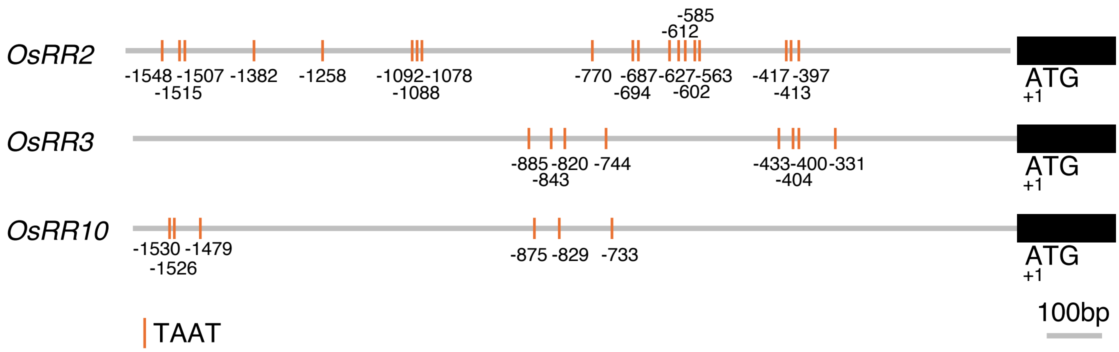


**Figure S6.** Positions of the putative binding sequence of TAB1 within the 1.6-kb upstream region of the *OsRR2*, *OsRR3*, and *OsRR10* genes. The putative TAB1 binding sequence 'TAAT' is indicated by the orange bars.

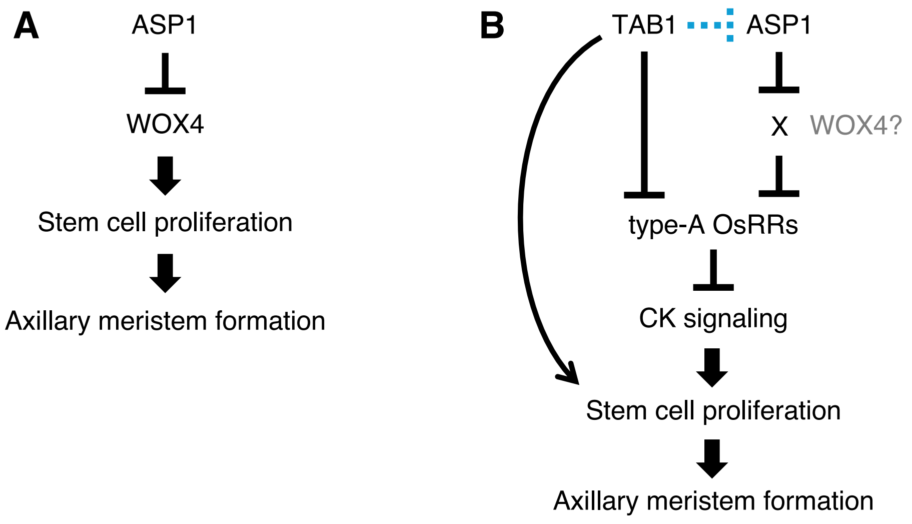


**Figure S7. Representation of the roles of TAB1 and ASP1 during axillary meristem formation. A)** ASP1 negatively regulates stem cell proliferation by repressing the precocious expression of *WOX4*. **B)** TAB1 promotes stem cell proliferation by activating cytokinin signaling through the downregulation of type-A *OsRR* genes. In contrast, ASP1 negatively regulates stem cell proliferation by promoting the expression of *OsRR* genes through an unknown factor 'X'. TAB1 may repress ASP1 function via posttranslational regulation, or it may act through an ASP1-independent pathway.
